# Automated workflow for the cell cycle analysis of (non-)adherent cells using a machine learning approach

**DOI:** 10.1101/2023.12.21.572803

**Authors:** Kourosh Hayatigolkhatmi, Chiara Soriani, Emanuel Soda, Elena Ceccacci, Oualid El Menna, Sebastiano Peri, Ivan Negrelli, Giacomo Bertolini, Gian Martino Franchi, Roberta Carbone, Saverio Minucci, Simona Rodighiero

## Abstract

Understanding the details of the cell cycle at the level of individual cells is critical for both cellular biology and cancer research. While existing methods using specific fluorescent markers have advanced our ability to study the cell cycle in cells that adhere to surfaces, there is a clear gap when it comes to non-adherent cells. In this study, we combine a specialized surface to improve cell attachment, the genetically-encoded FUCCI(CA)2 sensor, an automated image processing and analysis pipeline, and a custom machine-learning algorithm. This combined approach allowed us to precisely measure the duration of different cell cycle phases in non-adherent cells.

Our method provided detailed information from hundreds of cells under different experimental conditions in a fully automated manner. We validated this approach in two different Acute Myeloid Leukemia (AML) cell lines, NB4 and Kasumi-1, which have unique cell cycle characteristics. Additionally, we tested the impact of drugs affecting the cell cycle in NB4 cells. Importantly, our cell cycle analysis system is freely available and has also been validated for use with adherent cells.

In summary, this report introduces a comprehensive, automated method for studying the cell cycle in both adherent and non-adherent cells, offering a valuable tool for cancer research and drug development.

## Introduction

Cell cycle dynamics coordinate cellular division and proliferation through regulating the different cell cycle phases. Dysregulation in these processes is a hallmark of malignancies such as human cancer, where aberrant activities in cyclin-dependent kinases (CDKs), cyclins, and CDK inhibitors often drive uncontrolled proliferation. Consequently, targeting cell cycle components has emerged as a pivotal therapeutic strategy, especially crucial in pre-clinical drug evaluation (Malumbres and Barbacid, 2009; Khan and Wang, 2022).

Traditional methods for assessing cell cycle dynamics have been largely dependent on the quantification of DNA content through flow or image cytometry, providing a static snapshot of cell populations in various cycle phases (Furia, Pelicci and Faretta, 2013; Ligasová, Frydrych and Koberna, 2023). Although valuable, these techniques fall short in capturing intra-population variability and require additional protein markers for precise phase determination (Ligasová, Frydrych and Koberna, 2023; Rieger, 2022).

Methods utilizing cells expressing fluorescently labeled reporters and time-lapse microscopy can discriminate cell cycle phases at the level of individual cells, thereby offering valuable insights into the variability of cell cycle and cell cycle phase durations within the overall cell population (Chao *et al*., 2019; Hiratsuka and Komatsu, 2019).

Advanced imaging methods, such as time-lapse microscopy coupled with fluorescently tagged reporters, have shown promise in detailing cell cycle dynamics at the single-cell level. Technologies like the Fluorescent Ubiquitination-based Cell Cycle Indicator (FUCCI) have been employed for this, effectively demarcating cell cycle phases through color-coding (Sakaue-Sawano *et al*., 2017). FUCCI(CA)2 express the hCdt1(1/100) fused to mCherry fluorescent protein and hGem(1/110) fused to mVenus, generating a clear and distinct tricolor demarcation, separating G1 (red), S (green), and G2/M phases (yellow) (Sakaue-Sawano *et al*., 2017).

This dynamic insight is particularly crucial in the context of acute myeloid leukemia (AML), where chromosomal translocations generate fusion genes that disrupt cellular differentiation programs and drive proliferation (Alcalay *et al*., 2001).

The described experimental setup utilizes nanostructured titanium oxide-coated multiwell plates relying on the technology used in the commercially available Smart BioSurface (SBS) slides (Krol *et al*., 2021). Such technology should hypothetically enable us to immobilize non-adherent cells for extended imaging durations. To overcome the limitations of manual data analysis, we introduce an automated image analysis pipeline for time-lapse movies of AML cell lines, exploiting a FUCCI-based probe for visualization. Our data analysis approach combines custom image processing, TrackMate-based cell tracking, and machine learning-based track filtering. Thereby automating the entire data analysis workflow.

In summary, we present a comprehensive, experimental protocol for cell cycle analysis in adherent and non-adherent cells (summarized in **BOX 1**). The approach leverages routine imaging technologies and advanced data analyses, enhancing the precision and efficiency of drug screening protocols in oncological research.

## Results

### Modified conditions enable AML cells to adhere to the substrate feasible for live cell imaging

Live imaging and tracking of non-adherent cells, when multiple positions should be acquired, is challenging due to their high propensity to mechanical perturbations.

To conduct long time imaging of AML cells, we exploit the combined action of the SBS and the partial immobilization effect of methylcellulose (MC). We were able to image and track AML cells up to 72h when 20% complete medium was added to 80% MC and applied on AML cells previously adhered to the SBS (movie S1).

Two different AML cell lines, NB4 and Kasumi-1, were equipped with the FUCCI(CA)2 technology and chosen as the study models. NB4 cells have relatively faster doubling time in comparison to Kasumi-1 cells (Skopek *et al*., 2023). Hence, NB4 cells were treated with vehicle or CDK inhibitors, being compared to the naturally slow-cycling counterparts, Kasumi-1 cells, by time-lapse imaging (Fig. 1A and Fig. S1). As a result, we were able to follow single cells and the manual annotation of cell cycle phases according to the color of the red and green merged images in Fig. 1A, qualitatively confirmed the impact of CDK inhibitors. Furthermore, the expected difference in cell cycle progression of NB4 vs Kasumi-1 cells was evident.

**Figure 1:**
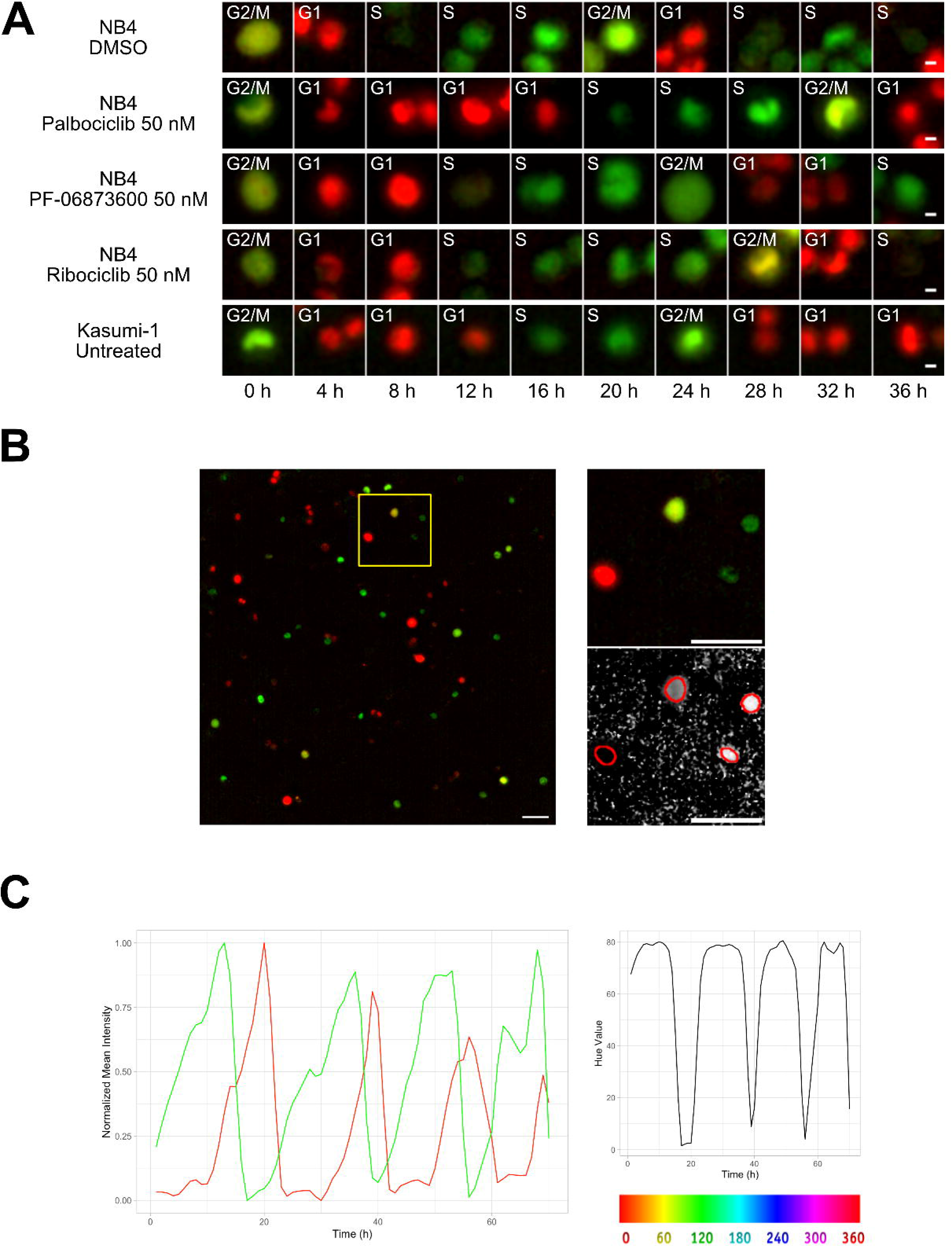
Cell Tracking. A) Examples from Kasumi-1 cell and NB4 cell lines (upon the different conditions: DMSO, Palbociclib 50 nM, PF-0606873600 50 nM, Ribociclib 50 nM, Untreated), showing a tracked cell that explores the different cell-cycle phases (Scale bar is 5 µm). B) Fluorescence images with R and G channels were processed in Fiji to create an HSB stack, from which the Brightness and Hue channel were extracted to be used respectively as tracking channel and cell phase identification channel (Scale bar is 50 µm). C) Plot curves generated using the data extracted from the TrackMate script execution. The variations over time of mCherry (red curve) and mVenus (green curve) fluorescence intensities, normalized between minimum and maximum values (left plot), and of the Hue scale (right plot) of a single NB4 cell in DMSO condition are shown.

### Image processing facilitates the cell tracking and profiling of cell cycle phases

The original time-lapse images were composed by Red and Green channels, detecting respectively the mCherry and mVenus markers of the FUCCI(CA)2 indicator. During the experiment each cell alternatively switches between the expression of these two markers as it goes through the different cell cycle phases, causing the lack of a single fluorescence channel suitable for tracking. Furthermore, the automatic profiling of cell cycle phases becomes cumbersome when dealing with two distinct channels that in the end give rise to two independent fluorescence time-series. We set up an image processing pipeline utilizing the open-source software Fiji (Schindelin *et al*., 2012) (NIH, version 2.14.0/1.54f) to transform the original images into a dataset optimized for the subsequent steps of tracking and cell cycle phase assignment. We represented the color changes that occur during the cell cycle with the Hue scale, as described in (Fujimoto *et al*., 2020). To achieve this, as represented in Fig. 1B, stacks of Red and Green channels were overlaid and converted into an RGB stack and then transformed into an HSB stack (Hue, Saturation and Brightness channels). The Hue channel was retained for the assignment of cell cycle phases, while the Brightness channel was used as tracking reference. These two were merged to the Red and Green fluorescence channels to form the final stack used for the tracking process.

In the tracking analysis, the Fiji plugin TrackMate (Tinevez *et al*., 2017) was employed, adapting the example script available on the dedicated TrackMate website (https://imagej.net/plugins/trackmate/scripting/scripting), to ensure the automation of the tracking step. The related parameters were selected and tuned according to each experiment, as well as the proper filters to discard uninformative tracks. The output consisted of a table with cells associated to selected tracks and their corresponding numerical features in each timeframe. Key features included the mean fluorescence intensity in the Red and Green channels, as well as the mean intensity of the Hue value.

The whole image analysis pipeline successfully recovered a unique tracking channel and effectively mapped the alternating red and green curves (Fig. 1C, left panel) into a single time series, as depicted in Fig. 1C, right panel.

### Incorrect track filtering by the machine learning model efficiently automates data cleaning and cell cycle phase assignment steps

The files generated by the Trackmate pipeline were imported into R (R Core Team, 2021) for the demultiplexing, filtering and data analysis. To address missing frames, a data imputation process was implemented, aimed at recovering instances with up to 5 consecutive missing values. This data imputation was executed using an Exponential Weighted Moving Average (EWMA) technique, employing a window size of 3, thereby encompassing 6 observations (3 preceding, 3 succeeding).

A smoothing spline approach was individually applied to the green, red, and hue channels, utilizing a regularization parameter (lambda) of 0.0001. A min-max scaling to transform value into the range [0, 1] was applied to make the fluorescence intensity and hue scale comparable among channels and samples. For the creation of the random forest model, a training pipeline was set up, as described in Fig. 2A. In accordance with Fig. 2B, feature extraction was carried out on the green and red channels to derive temporal descriptors of the data.

**Figure 2:**
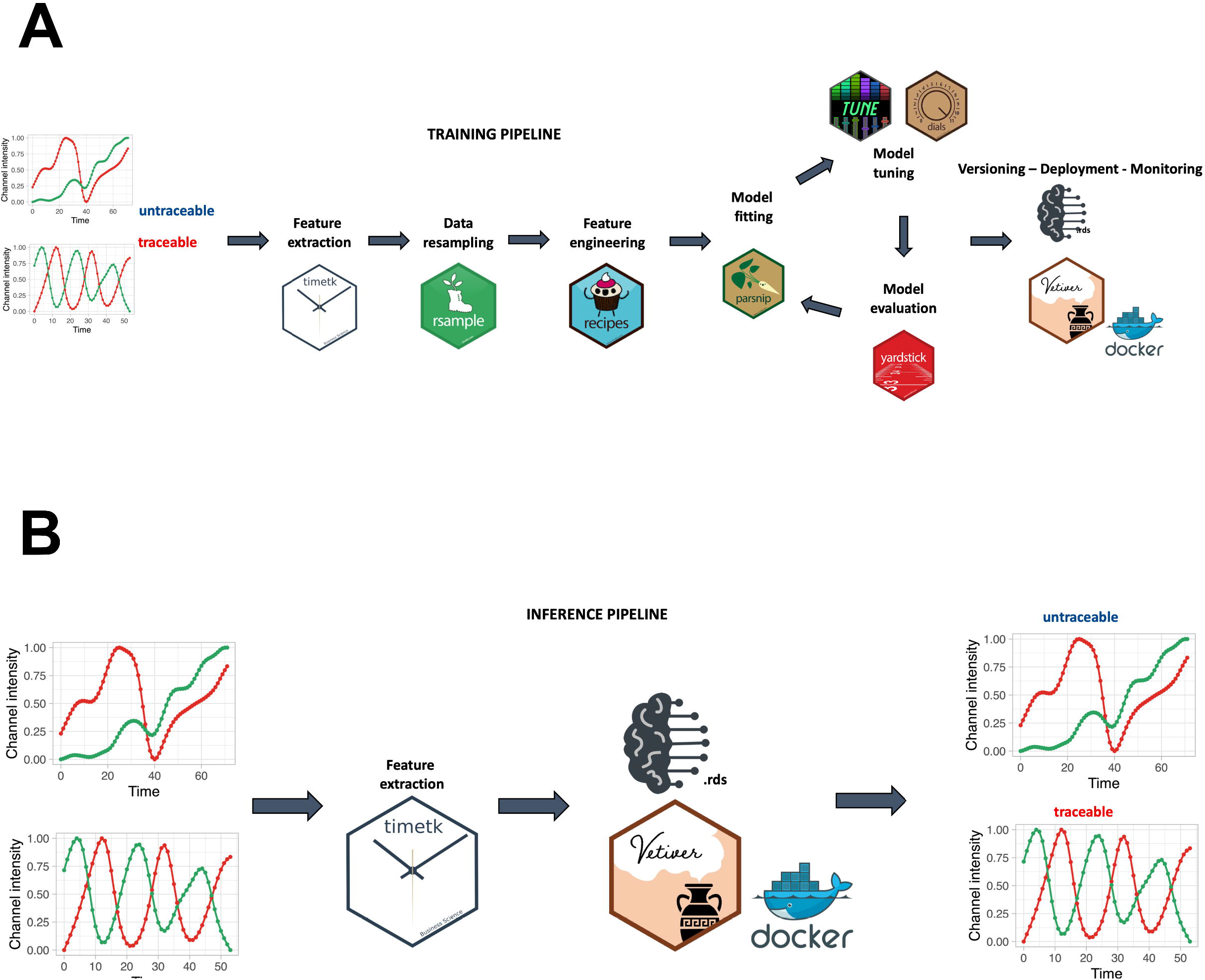
Training and Inference Pipelines. A) The Machine Learning pipeline followed to create the quality model. Using timetk time-series associated features are extracted from the list of manually annotated tracks. A random forest model is then trained to predict whether a track is cycling or not. B) An unannotated track can be fed to the model to predict whether it is cycling or not.

Employing these extracted features, the trained random forest model was applied to discriminate between trackable and untraceable cells. Subsequently, only the cells predicted as traceable by the model were retained for subsequent analyses. Here are reported the obtained number of cells for each condition, merging all the experiments performed:

- NB4 DMSO: 410 cells (3 experiments) out of 1862 tracked cells
- NB4 Palbociclib 50 nM: 328 cells (3 experiments) out of 2881 tracked cells
- NB4 PF-06873600 50 nM: 206 cells (1 experiment) out of 1102 tracked cells
- NB4 Ribociclib 50nM: 119 cells (1 experiment) out of 784 tracked cells
- Kasumi-1 Untreated: 119 cells (1 experiment) out of 1604 tracked cells
- MDA-MB-231 Untreated: 1116 (1 experiment) out of 3204 tracked cells

By using the hue channel each track was partitioned into its cell cycle component using the following thresholds:

• **G1**: hue >= 0 & hue < 0.65

• **S**: hue >= 0.85

• **G2/M**: hue >= 0.65 & hue <= 0.85

Additionally, a refinement step involving cell reassignment was conducted to identify instances where the G1 to S phase transition was labeled as G2/M in frames preceding the S phase. The outcome of the analysis is visually depicted in Fig. 3A and B.

**Figure 3:**
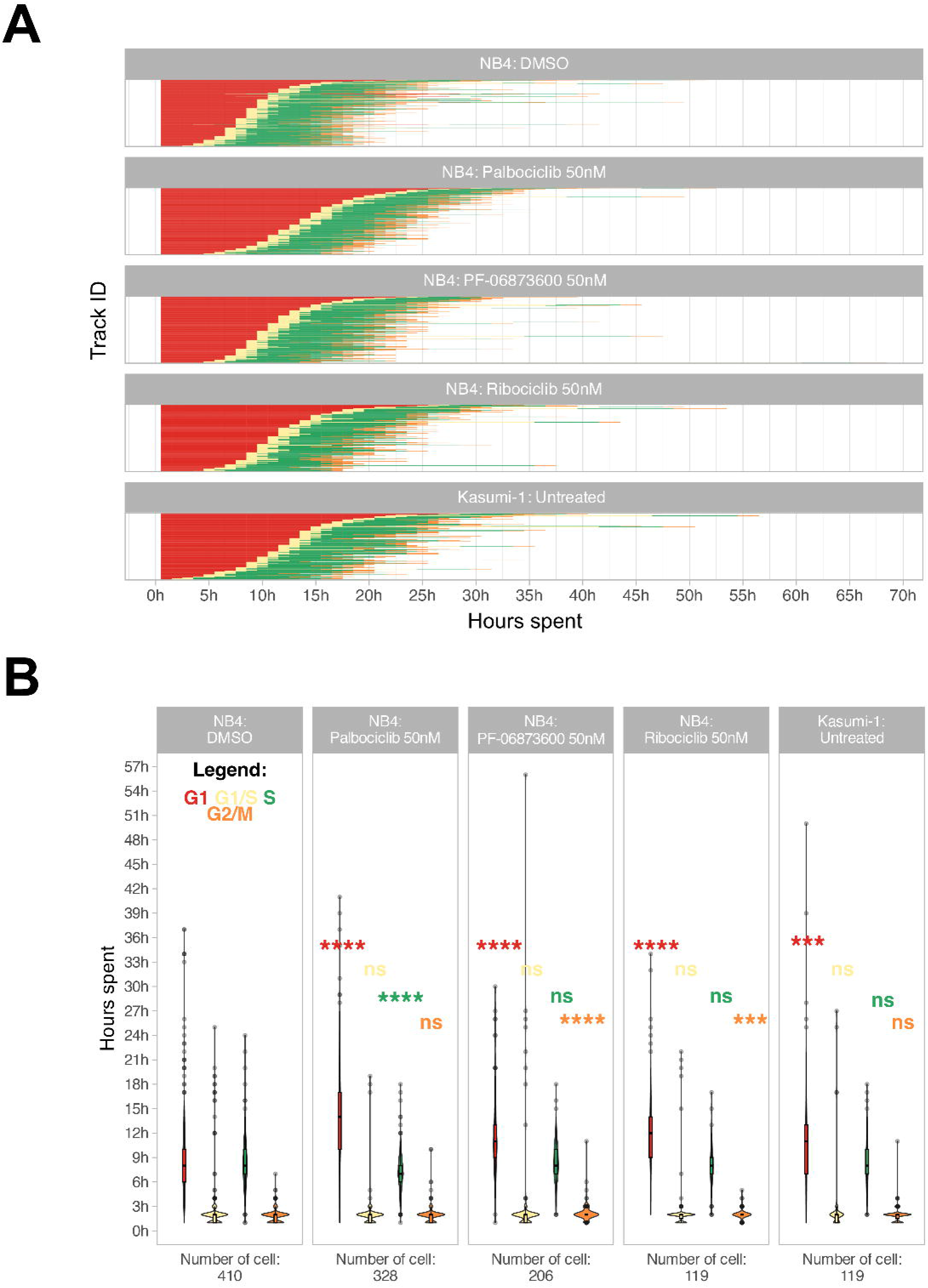
Cell Cycle Phase Assignment. A) Waterfall plot of treated and untreated AML cells. Each row corresponds to a single cell. B) Boxplot of the cell phase duration of the first cell cycle, as obtained by the pipeline, without excluding of the outliers. The asterisks are the adjusted significance level given by Wilcox test of the sample for each over the NB4 DMSO for each phase according to the color (ns: p > 0.05, *: p <= 0.05, **: p <= 0.01, ***: p <= 0.001, ****: p <= 0.0001).

### Evaluation of tracks’ selection performances confirms the efficiency of the machine learning model

We carried out a validation of the performances of the whole pipeline in selecting “traceable tracks. Specifically, we extracted from one experiment the ID of the tracks that belonged to the vehicle (DMSO) treated condition identified as “traceable” by the ML algorithm. We then validated these tracks by visual inspection of the corresponding images, checking for eventual tracker errors We created a scoring metric defined as follows:

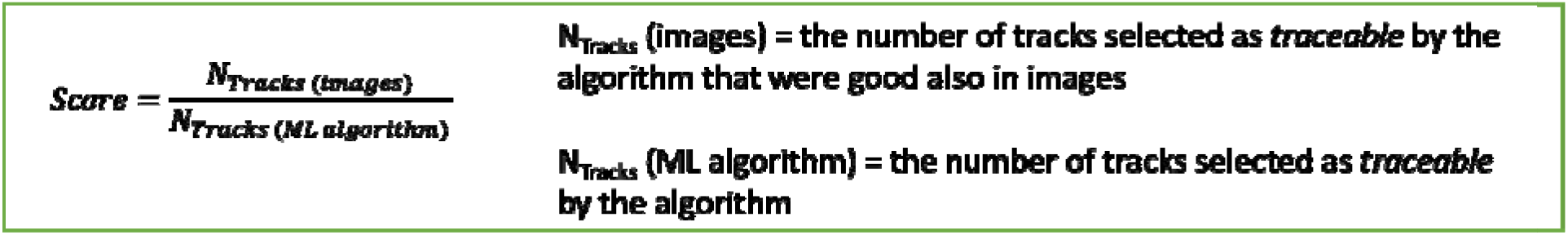

We checked up to 102 tracks that underwent to whole cell cycle profiling by the analysis pipeline and we found that, among these, 63 were also consistent with images (Score = 0.62). We then decided to select tracks identified as traceable with a probability greater than 0.70, gaining a final Score of 0.74 (57 tracks consistent with images over 77 tracks).

### The workflow allows the identification of the G1 to S phase transition

The empirical observation of the superimposition of red and green fluorescence signals during the G1 to S transition makes the classification of the cell in one of the two dual states challenging. Cells often exhibit a brief gap in fluorescence during this transition, as illustrated in the first image categorized as S phase in both Fig. 1A and Fig. S1. This apparent lack of fluorescence occurs because mCherry fluorescence rapidly decreases to near zero at the start of the S phase, while mVenus fluorescence begins to increase slowly (Fig. 1C). Although seemingly non-fluorescent, this initial S phase is marked by the coexistence of both mCherry and mVenus fluorescence, producing a yellow hue in the scale before the S phase, labeled as the G1 to S transition (G1/S) by the refinement step in our model. Due to the lower intensities of mCherry and mVenus during this transitional phase, compared to other cell cycle phases, the yellow fluorescence is barely visible in fluorescence images where brightness and contrast settings are dominated by the bright G1 red and S green fluorescence.

To evaluate the relative expression of the two fluorescent probes during this transition, we conducted flow cytometry and confocal experiments on Kasumi-1 and MDA-MB-231 cells expressing the FUCCI(CA)2 probes. In these experiments the S phase was labeled by a pulse of EdU and the DNA by DAPI or Hoechst® 33342 (Fig. 4 and Fig. S5). Fig. 4A shows cell cycle profiling based on EdU vs. DAPI can successfully discriminate G1, S, and G2/M phases, with the S phase being subdivided into early, mid, and late S phases (Fig. 4A and Fig. S5A). The proportion of cells in each phase is comparable to the cell cycle profiling of the same cells by FUCCI(CA)2 (Fig. 4B and Fig. S5B). Moreover, the temporal transition through the S phase discovered by EdU correlates with the mVenus emission levels detected by FUCCI(CA)2 alone (Fig. 4B and Fig. S5B). This is evident by the accumulation of the EdU-early S phase cells on the left side of FUCCI(CA)2-S phase cells (low mVenus emission). On the contrary, the EdU-late S phase cells tend to accumulate on the right side of the FUCCI(CA)2-S phase cells (high mVenus emission) (Fig. 4B and Fig. S5B). This finding is in line with the earlier observation of the relatively rapid loss of mCherry at late G1 and the gradual increase in mVenus emission throughout the S phase (Fig. 1C). These observations suggest the possibility that FUCCI(CA)2 alone can discriminate early vs. late S phase cells. However, this aspect can be exploited in cell populations with necessarily similar levels of FUCCI(CA)2 expression.

**Figure 4:**
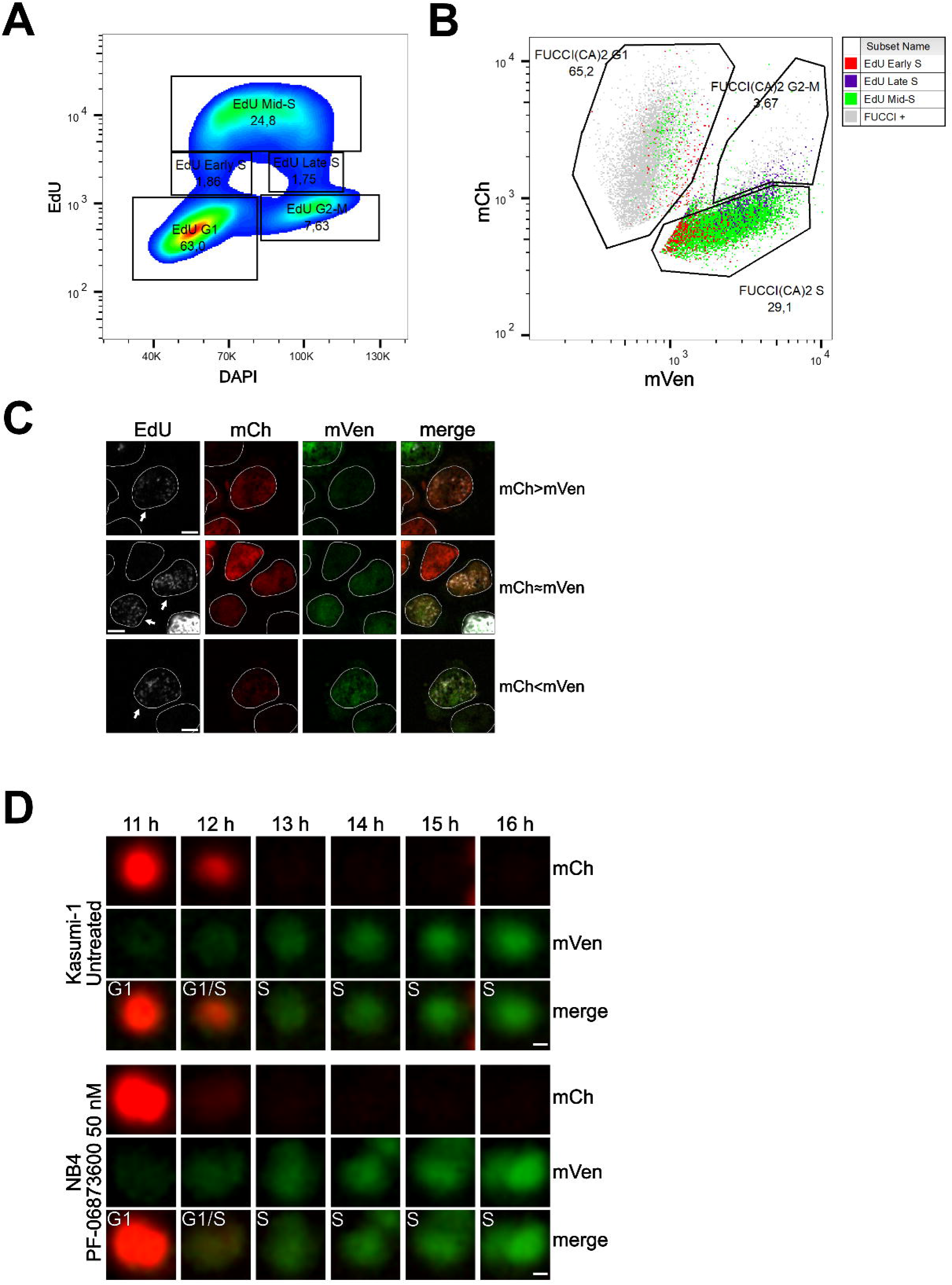
Cell Cycle Profiling and G1 to S Transition Analysis. A) Density plot showing the cell cycle profiling of FUCCI(CA)2-Kasumi-1 cells based on EdU vs DAPI. B) Scatter plot of the same cells as in “A” showing the cell cycle profiling based on FUCCI(CA)2. Cells identified to be in early, mid, and late S phase by EdU are highlighted in red, green, and purple, respectively. Due to the chosen gates and the 2-hour EdU pulse, a small percentage of EdU-positive cells leaked into the G1 and G2-M FUCCI(CA)2-defined groups. C) Representative images of nuclei during the G1 to S transition are indicated by arrows. Depending on the exact time after S phase initiation, the overlay of the mCherry and mVenus channels can show a red prevalence (first row), a yellow color (middle row), or a green prevalence (third row). The scale bar represents 5 µm. D) The frames, including the transition from G1 to S phase of the untreated Kasumi-1 cell and the NB4 PF-06873600 treated cell shown in Fig. 1A, are presented from the 11-hour to the 16-hour frame. In these image sequences, the G1/S label was assigned to the first frame showing dim but detectable mVenus fluorescence and dim but still detectable mCherry fluorescence. The scale bar represents 2 µm.

Interestingly, from Fig. 4B is evident that the EdU-early S phase cells can express mCherry and mVenus at different levels. Fig. 4C shows segmented nuclei of cells at the initial S phase (arrows), characterized by dim EdU staining and DNA amount only slightly higher than 2N (not shown), where the dim green and red fluorescence overlaid determined a predominant red (when the intensity of mCherry is slightly higher than the intensity of mVenus), yellow (when the intensity of mCherry and mVenus are similar), or predominant green (when the intensity of mVenus is slightly higher than the intensity of mCherry) color in the merged image. The same phenomenon was observed in MDA-MB-231 cells (Fig. S5). We examined this phenomenon in three of the previously shown cells - Kasumi-1 and PF-06873600-treated NB4 in Fig. 1A and MDA-MB-231 in Fig. S4C - where all the available frames were visually evaluated. The G1 to S transition was once captured as predominantly red (mCherry > mVenus, Fig. 4D), once as yellow (mCherry ≈ mVenus, Fig. 4D) and once as green (mVenus > mCherry; Fig. S5D).

### The method can quantify the cell cycle progression and does not impact the transcriptome over long time periods

We were able to accurately quantify the cell cycle progression of NB4 and Kasumi-1 cell lines over 72 hours at single-cell resolution and, as represented in Fig. 3A and B, cell cycle differences of NB4 and Kasumi-1 cells were quantified. Moreover, the method was able to discriminate cell cycle differences of leukemic cells in presence of various sub-optimal doses of different CDK inhibitors. These CDK inhibitors are potent inhibitors of CDKs2/4/6, which mainly regulate the G1 to S transition (Fassl, Geng and Sicinski, 2022). Hence, we expected the administration of nanomolar concentrations of these agents to lengthen mainly the G1 phase with less impact on the other phases. Our live cell imaging of the relatively fast-cycling NB4 cells in presence of these drugs at such spectrum of concentrations affirmed the significant G1-prolongation in these cells (Fig. 3B and movies S2-4). The total average duration of one full cell cycle was quantified as 21.5 h ± 6.5 h (mean ± standard deviation) for NB4 cells and 24.0 h ± 7.8 h for Kasumi-1 cells. For instance, administration of 50nM Palbociclib extended mainly the G1 phase of the NB4 cells by about 5 hours (from 9.1 h ± 5.1 h to 14.2 h ± 5.7 h in vehicle treated and Palbociclib treated NB4 cells, respectively).

It was demonstrated that the nanostructured surfaces can promote changes in the cellular protein expression profile (Schulte et al., 2016). We thus questioned the possible impact of the SBS on gene expression profile of the AML cells. We made a comparison of transcriptome of the cells pre- and post-imaging by performing RNA-seq investigations. As represented in the supplementary Fig. S2, we did not observe any meaningful transcriptomic alterations throughout the process. Hence, we conclude that the protocol is feasible for biological investigations without a significant effect on the transcriptomic profile of the cells.

## Discussion

In the last few years, efforts have been made to simplify the cell cycle assessment relying on FUCCI technology, resulting in the development and public availability of software tools and ImageJ plugins (Roccio et al., 2013; Koh et al., 2017; Ghannoum et al., 2021; Taïeb et al., 2022). However, all these methods require human intervention at different points in the analysis workflow, such as the initial selection of cells for analysis (Taïeb et al., 2022) or the manual correction of inaccurate tracks during the analysis (Roccio et al., 2013; Koh et al., 2017; Ghannoum et al., 2021; Taïeb et al., 2022). In this report we describe a complete protocol for the cell cycle analysis of adherent and non-adherent cells expressing the FUCCI(CA)2 technology in a fully automated manner. The complete experimental workflow is explained in BOX1 and illustrated in Fig. 5. It was applied to the analysis of the cell cycle phases of two different AML cell lines, NB4 and Kasumi-1, which have different durations. Hence, detecting this difference was a reliable initial verification strategy for the protocol’s efficiency. As a second quality check step, we decided to treat the relatively fast-cycling NB4 cells with suboptimal concentrations of three different CDK inhibitors (Palbociclib, PF-06873600 and Ribociclib). These CDK inhibitors have different inhibitory impact on various CDKs (Freeman-Cook et al., 2021; Fassl, Geng and Sicinski, 2022). The effect of the three CDK inhibitors on the cell cycle duration of NB4 cells was evaluated. Up to about 400 cells in one single experimental condition were quantified, for up to 12 different conditions in a single experiment. This entire analysis process took approximately 2 hours of human involvement, for a total execution time that ranges from 12 to 48 hours, depending on dataset size, that is commissioned to a machine (Fig. 4). It should be taken into consideration that the manual correction of incorrect tracks takes approximately 2 hours per field of view (FOV). Hence, in the reported experiments of this study, the analysis of a single well would take about 40 to 50 hours of manual work. This amount of time would make such analysis unfeasible. Not only the suggested protocol makes these experiments doable, but also the experimental set-up can be scaled up according to the experimental needs. We successfully emphasized alterations of few hours in the duration of cell cycle phases when NB4 cells were subjected to exceedingly low concentrations of inhibitors. Moreover, the set-up pipeline makes possible the quantification of potentially thousands of cells in an automated fashion relying on the crucial contribution of a machine learning algorithm.

**Figure 5:**
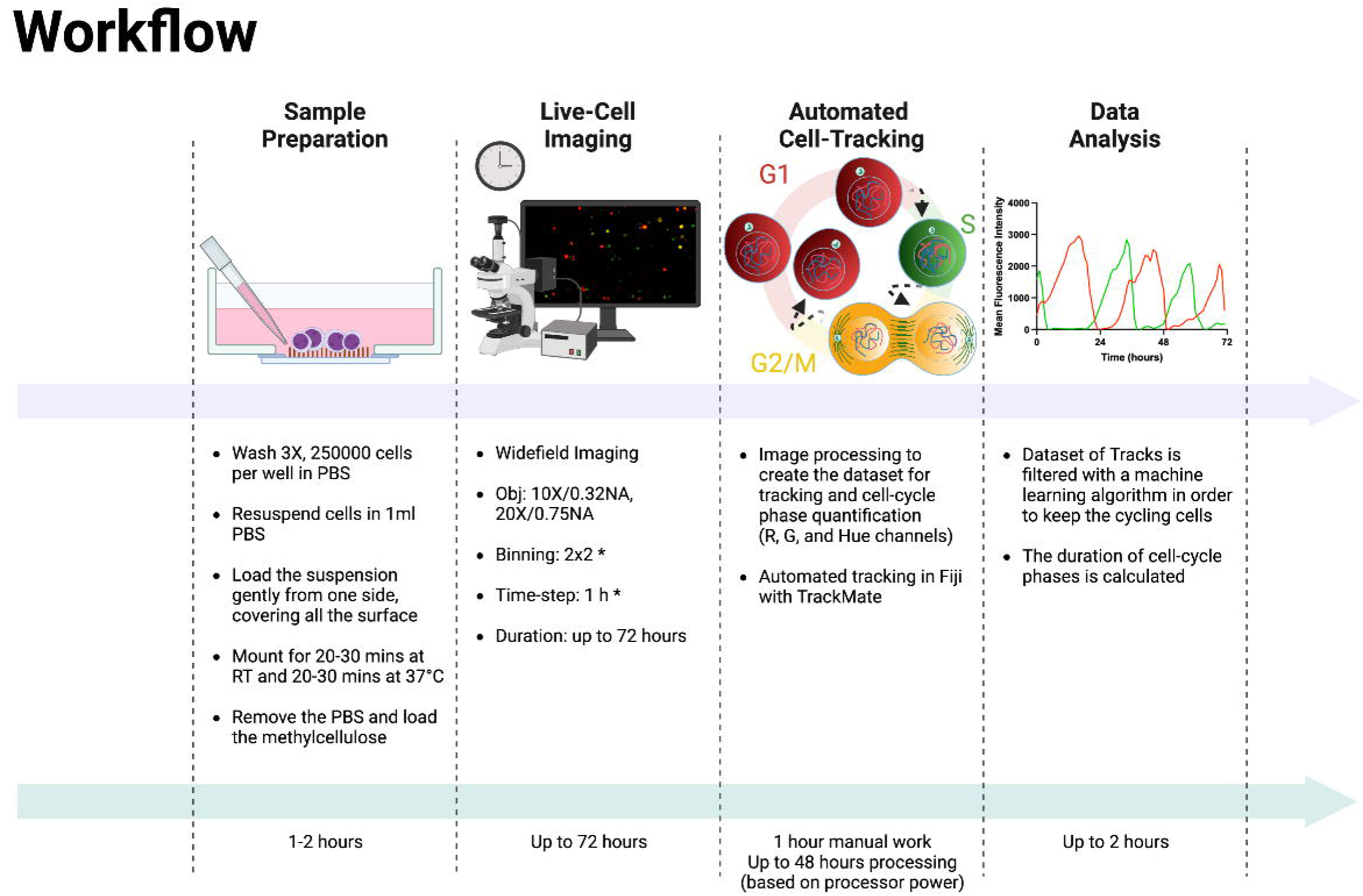
Experimental and Analysis Workflow. Summary of the entire experimental workflow, from cell seeding to the calculation of cell cycle phases. Asterisks in the Live Cell Imaging section mean optional settings.

Various approaches have been documented for imaging non-adherent cells. These range from creating custom supports to confine cell movement within a restricted region enabling extended imaging periods of up to 40 hours (Day et al., 2009), to partially immobilizing cells using substances like gelatin (Ritter et al., 2020) or low-melting-point agarose (Strong and Daniels, 2017).

The capability to track non-adherent cells for up to 72 hours (almost equivalent to three complete cell cycles in NB4 cells) was achieved by employing SBS-coated glass in combination with the addition of MC to the culture medium. MC was necessary to limit the movement of single cells or cell clusters after more than 24 hours of observation (movies S1 and S5). We deduce that this seeding protocol has the potential to be effectively employed with various non-adherent cell types. This is based on the principle that cell adhesion on SBS is facilitated by the nanostructure’s ability to engage with integrins (Schulte et al., 2016), which are widely expressed in diverse cell types (Johansen et al., 2018; Floren et al., 2020; Kim et al., 2020; Ogana et al., 2023). The seeding protocol could also certainly be extended to other types of live cell imaging experiments, especially when the fluorescence intensity can be monitored for shorter durations, thereby circumventing the need for the addition of MC. Moreover, the image and data analysis pipelines can be easily applied to adherent cells, as we showed in Fig. S4.

The presented image and data analysis workflows rely on 2 different software, Fiji and R, widely used by the imaging and data analysis community. The scripts were made publicly available, enabling customization to tailor the workflow according to specific requirements. This not only streamlines the cell cycle analysis of cells expressing the FUCCI(CA)2 indicator, but also makes it accessible to researchers with basic programming knowledge.

In conclusion we confirm the efficiency of the optimized conditions in immobilizing the non-adherent cells for long time periods, without affecting their predicted response upon different environmental circumstances. The described work allows researchers in the field to analyze thousands of (non-)adherent cells per each experimental condition, by using the image and data analysis pipelines to automatically perform image processing, cell tracking, filtering of incorrect tracks and cell cycle phases identification and characterization.

### BOX 1 – experimental and analysis workflow

Sample Preparation (in-suspension cells): The cells of interest should be at optimum viability and physiological conditions prior to the mounting process on the SBS-multiwell plates. This step requires 1-2 hours of manual handling and is divided into seven steps as follows: (I) Consider 250 thousand cells for one well of a standard 12-well SBS-multiwell. Wash the cells 3 times in sterile phosphate-buffered saline (PBS). Note 1: PBS should be at room temperature (RT). Note 2: The media and PBS should not be vacuumed during the process. It is crucial to make sure the cells won’t get dried at any point of the process. Note 3: The mentioned number of cells are for cells with an average size of 10 to 20 micrometers in diameter and a doubling time of 20-30 hours. The number of cells with different properties, should be optimized accordingly. (II) Resuspend the cells in 1000 µl PBS and gently mix, achieving a homogenous cell suspension. (III) Load the suspension slowly from one side of each well, covering all the surface. (IV) Let the cells mount for 20-30 mins at RT, followed by 20-30 mins at 37°C. Note 4: Avoid moving or shaking the plate or the surface underneath. (V) Remove the PBS with pipette (we suggest not to use the vacuum and not to tilt the well). (VI) Wash the wells 2 times with enough volume of media without fetal bovine serum (FBS) (we suggest not to use the vacuum and not to tilt the well). Note 5: Any protein contamination, including FBS can easily perturb adhesion of the cells on the titanium-oxide particles. Note 6: Load and aspirate slowly from the side of the wells, keeping the pipette at 45° angle. (VII) Load enough methylcellulose gently from one side of the well and proceed to the next step. Note 7: In case any treatments are needed during the time-lapse experiment, the desired agents to can be added to the methylcellulose compartment prior to loading on the wells. Sample Preparation (adherent and semi-adherent cells): In case the cells of interest are (semi-)adherent cells, plate the cells at optimum confluency (based on the planned time-lapse duration and cell properties) on glass-bottom dishes and proceed to the next step. Live-Cell Imaging (I) Pre-heat the microscope incubator at 37 °C and 5 % of CO2 for standard cell culture. Otherwise tune the temperature and humidity controller according to the ideal cell viability conditions. (II) Mount the sample on the microscope stage and choose the imaging parameters depending on the cell type. Note 8: we suggest setting the objective and image binning to represent each cell with at least 15 pixels in diameter. Note 9: we suggest adjusting the excitation intensity and exposure time for the detection of mCherry and mVenus signal to have a difference between maximum and minimum gray value of at least 1000 units (for 16-bit camera). (III) Select the desired total duration of the experiment and the time-frame parameter, considering the cell type viability upon illumination and the speed of cells movement. Note 10: for tracking purposes it is convenient to start with a minimum time interval of 30 min and evaluate the outcome of the time-lapse, adjusting eventually the described parameter in the following experiments Image processing (I) Open the pre-processing macro (Image_processing_HSB_v1.ijm) in your Fiji Note 11: the execution of the macro requires the prior installation of the Basic plugin (https://github.com/marrlab/BaSiC) for flat-field correction (II) Set the proper image filters and background subtraction processing for red and green channels at lines 58-78 and 85-94 (III) Choose the maximum values of the Brightness and Contrast for both red and green channels and set the found values at lines 116 (red) and 118 (green) (IV) Set the proper value of the Top Hat filter on the Brightness channel (HSB stack) at line 127 (V) Set the correct position of red and green channels, depending on the experiment, at lines 173-174. (VI) Run the macro from the Fiji script editor (VII) Set in the dialog window the input directory (where original data are stored), the output directory (where you want to store processed images) and the original file format

Tracking Analysis

I. Open the TrackMate script in the Fiji macro editor
II. Adjust the parameters settings (from line 39 to 58) according to the experiment
III. Run the script from the Fiji script editor
IV. Set in the dialog window the input directory (where the images processed with the previous pipeline are stored) and the output directory (where you want to store the results table)

Data Analysis

Sample/condition demultiplexing

I. Read the list of files containing tracking information for each cell. This can be done by looping through the files and reading into R using the function read.delim().
II. Merge all the tables into a single table using the filename to create a column for the demultiplexing of the condition.

Missing frame imputation

I. Use the na_ma() function to perform imputation. This function let the user choose among different weighting strategies, we suggest trying the “exponential” or the “simple”, as well as trying different values for k (dimension of the moving average window) and maxgap (maximum number of successive NA values).

Data smoothing

I. Use the function smooth.spline() to remove noise from the timeseries and enhance the trend. We suggest trying different values for lambda value starting with a very low one such as [1e-05, 1e-04, 1e-03].

Data normalization

I. Use the function normalize_vector() to make the curves comparable to each other.

Feature extraction

I. Use the function tk_tsfeatures () to extract features from the channel intensity, different features can be specified to be extracted such as:

a. The number of times the time series crosses its median.
b. Autocorrelation of the time series.
c. Autocorrelation of the first/second-differentiated time series.
d. Spectral entropy.
e. Stability and lumpiness on a tilled version of the timeseries.

Traceability assessment

I. Use the function predict() passing the table and the pre-trained model to predict whether a track is traceable or not. The output is a table containing the predicted class as well as the predicted probability of being part of a specific class.
II. The predicted probability can be used to further filter the predicted traceable cells.

Cell phase assignment

I. The cell cycle phase quantification is performed on the cells predicted as traceable; this is done by setting two thresholds on the Hue intensities. We suggest trying different values according to the original Hue intensity reached by the fluorescence observed.
II. The first round of phase assignment divides the tracks into:

a. G1
b. S
c. G2/M
III. By looping through each track, a second round of phase assignment is performed to re-assign frames wrongly classified as G2/M that occur before S. Three different strategies can be followed:

a. Re-assign the G2/M frames to S
b. Re-assign the G2/M frames to a new G1/S phase

Cell phase quantification

I. Group your data using the group_by() function using the Track_ID and Condition columns as grouping variables then use the function add_count() to count the number of frames in each condition for each track.
II. Divide the number obtained by the number of frames acquired in an hour to obtain the time spent in each phase (in hours units).
III. Sum the time spent in each phase to obtain the total cell cycle duration.

## Methods

### Key resource table

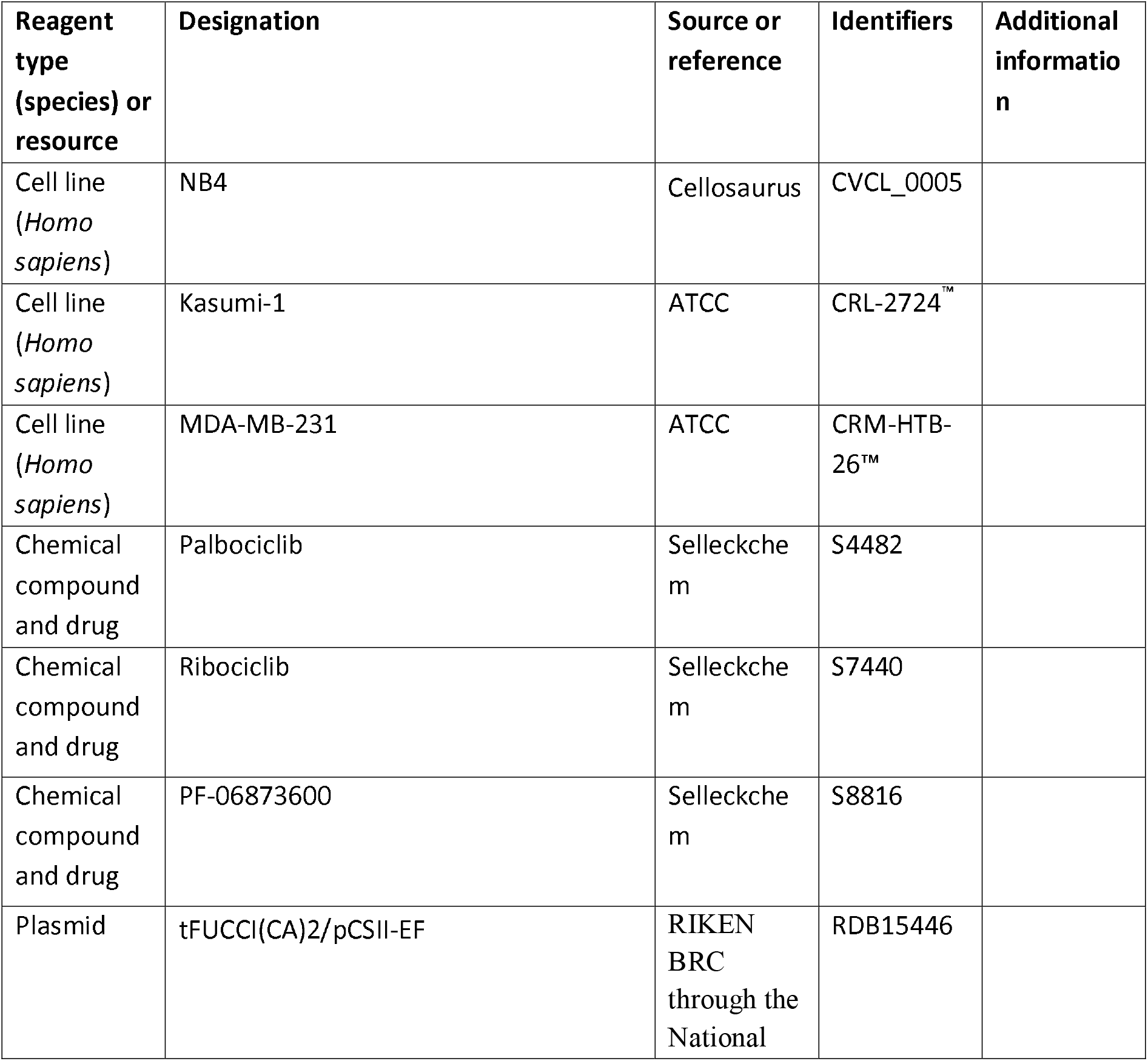
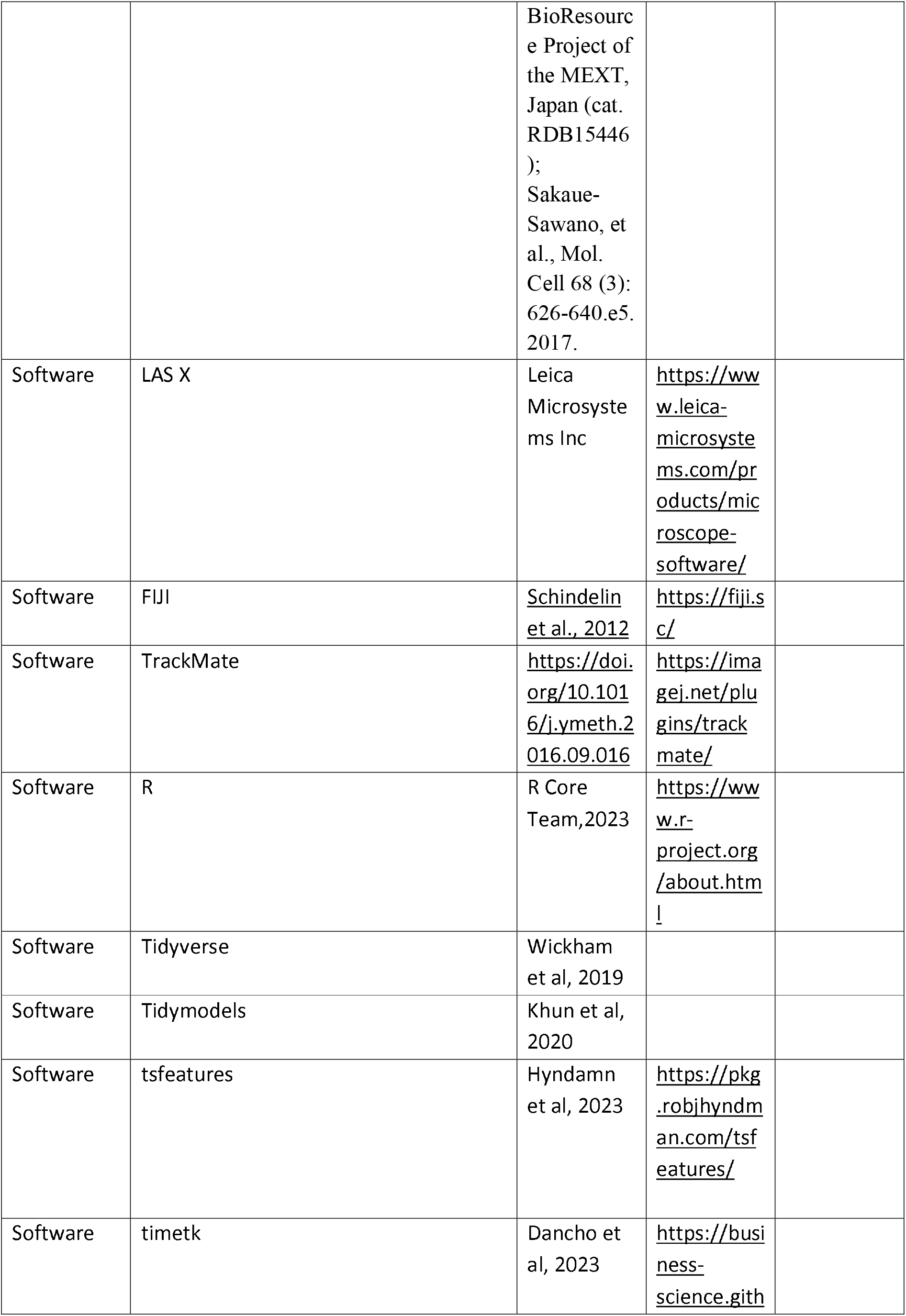
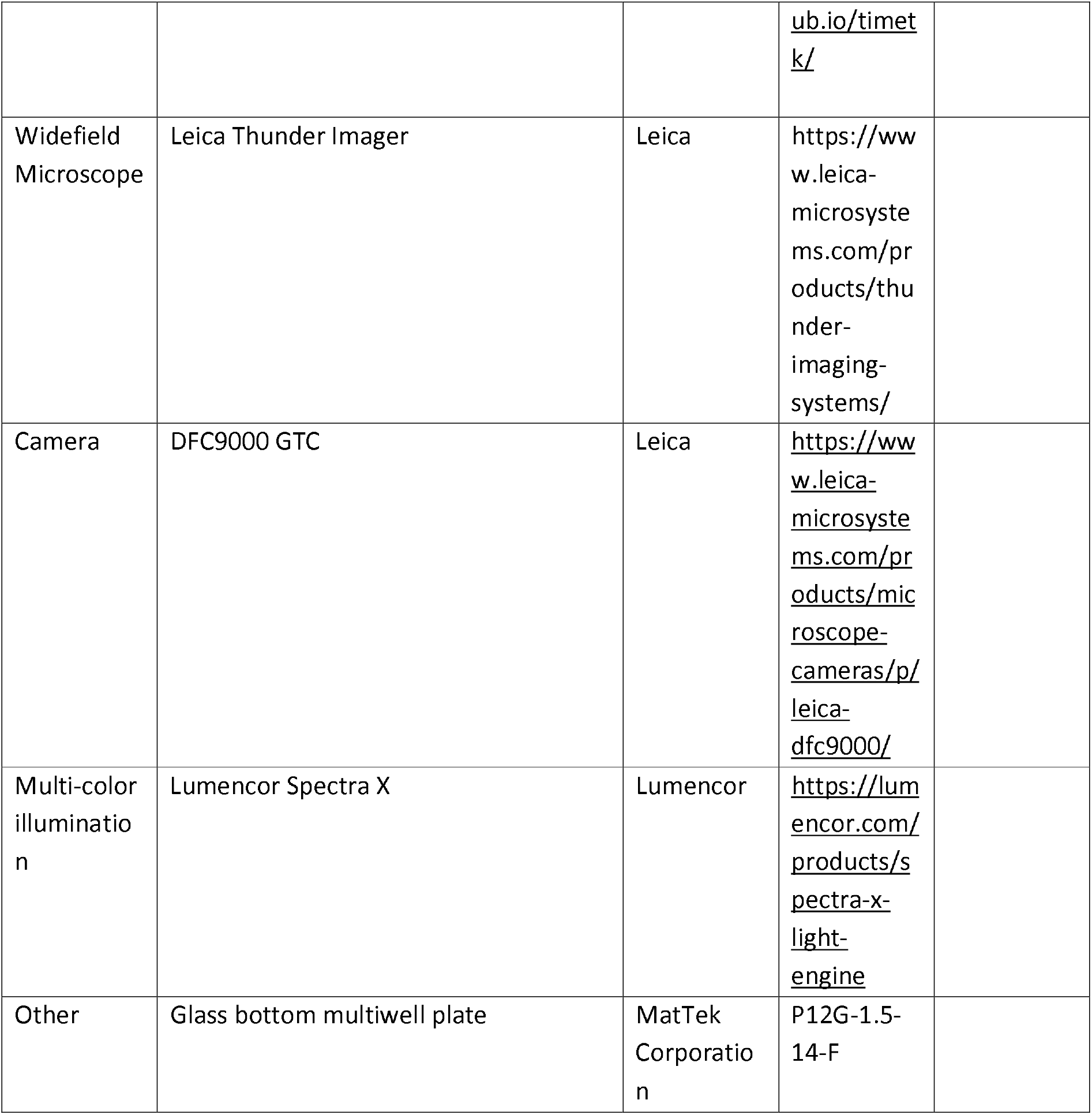

### Cell culture

#### Cell lines and growing conditions

NB4 and Kasumi-1 cells were grown in 3ml of a mix of 80ml MethoCult™ H4230 RPMI-1640 plus 20ml Roswell Park Memorial Institute (RPMI) 1640 medium with final concentrations of 10% fetal bovine serum (FBS), 2mM glutamine and 1% Penicillin/Streptomycin. MDA-MB-231 cells were grown in Dulbecco’s Modified Eagle Medium (DMEM) with 10% FBS+2mM L-Glutamine and 1% Penicillin/Streptomycin. All cells were grown according to ATCC recommendations in a humidified tissue culture incubator at 37 °C with 5% CO2 environment.

#### Generation of FUCCI(CA)2-equipped cell lines

Cells were put in 24 well-plates and plated at a density of 500,000 cells in 500 μl of medium per well. The Lentiviral vector carrying the tFUCCI(CA)2/pCSII-EF (Sakaue-Sawano *et al*., 2017) plasmid was diluted in RPMI 10% serum pen/strep in order to add 500 μl to the cells. Three rounds of infection were carried out in the presence of 5ug/mL of polybrene (Sigma). Cells were infected simply by adding the 100X concentrated virus supernatant (20-35μL/well) onto the cells; the plate was centrifuged for 1 hour at RT at 2500rpm. The medium is added 2 hours post-infection up to 1ml final. Then incubated at 37°C overnight. Infected cells expressed FUCCI(CA)2 probes, emitting mCherry and mVenus. Therefore, the selection of the cells expressing the FUCCI(CA)2 was done by sorting the fluorescence markers, mCherry and mVenus on fluorescence-activated cell sorting (FACS) instrument (FACSAria cell sorter,BD Biosciences, Oxford, UK).

### SBS coated bottom-glass multiwell plates production

Glass bottom multiwell plates are coated by Tethis using the Supersonic Cluster Beam Deposition (SCBD) technology (a detailed description of SCBD and its principle of operation can be found in Refs.(Piseri, Tafreshi and Milani, 2004; Wegner et al., 2006). A supersonic seeded beam of titania nano-clusters is produced, under high vacuum, by a pulsed microplasma cluster source (PMCS) (Piseri, Tafreshi and Milani, 2004; Wegner et al., 2006) and deposited on glass bottom multiwell plates. The process is tuned to produce nanostructured TiO2 films (thickness from 50 to 200 nm) with a controlled nanoscale morphology. The titania nanostructured coating of the plates is transparent and biocompatible and has a surface topography that promotes the spontaneous adhesion and immobilization of living cells (Carbone et al., 2006).

### Sample preparation

Approximately 250,000 of Kasumi-1 and NB4 cells and 100,000 of MDA-MB-231 cells were plated either in each well of the Tethis SBS 12-well multi-wells or 8-well ibidi plates, in triplicates in the presence of the aforementioned compounds at day 0 (T0). Cells were treated with final 2nM, 10nM, 50nM, 250nM, and 1250nM of Palbociclib, 50nM, 75nM, 100nM, 125nM and 150nM of Ribociclib, and 25nM, 50 nM, 75nM, 100nM and 125nM of PF-06873600, diluted in advance and directly in the relevant culture medium for each cell line. Only the 50nM treatments are shown for Ribociclib and PF-06873600.

Three cell suspension samples were prepared for triplicate independent counting and the average of three readings used as the cell count. Cell suspension was diluted with Trypan Blue dye (Sigma)^TM^ at 1:1 ratio to identify cell viability using Biorad TC20 automated cell counter.

### RNA extraction and RNA-seq protocol

Total RNA was extracted from dry pallets of cells collected prior- and post-acquisitions and purified using the Zymo Research Quick-RNA Miniprep (W/O directzol). Reverse transcription was performed with the SuperScript II Kit (Invitrogen), according to the manufacturer’s protocol. RNA-seq was performed according to the True-seq Low sample protocol selecting only polyadenylated transcripts. In brief, before starting mRNA isolation and library preparations the integrity of the total RNA was evaluated by running samples on a Bioanalyzer instrument by picoRNA Chip (Agilent), then converted into libraries of double stranded cDNA appropriate for next generation sequencing on the Illumina platform. The Illumina TruSeq v.2 RNA Sample Preparation Kit was used following manufacturer’s recommendations. Briefly, 0.1-1 μg of total RNA were subjected to two rounds of mRNA purification by denaturing and letting the RNA bind to Poly-T oligo-attached magnetic beads. Then fragmentation was performed exploiting divalent cations contained in the Illumina fragmentation buffer and high temperature. First and second strand cDNA is reverse transcribed from fragmented RNA using random hexamers. First strand cDNA was synthesized by SuperScript II (Invitrogen) reverse transcriptase and random primers and second strand cDNA synthesized by DNA polymerase I and Rnase H. The subsequent isolation of the cDNA was achieved by using AMPure XP beads (depending on the concentration used, these beads can efficiently recover PCR products of different sizes). The product recovered contained overhanging strands of various lengths due to the fragmentation procedure. The 5’ and 3’ ends of cDNA are repaired by the 3’-5’ exonuclease activity and the polymerase activity and adenylated at 3’ extremities before ligating specific Illumina oligonucleotides adapters followed by 15 cycles of PCR reaction using proprietary Illumina primers mix to enrich the DNA fragments. Prepared libraries were quality checked and quantified using Agilent high sensitivity DNA assay on a Bioanalizer 2100 instrument (Agilent Technologies).

### RNA sequencing data analysis

Raw reads 51bp PE for NB4 and Kasumi-1 cells were quality-filtered and aligned to the hg18 reference genome using nf-core/rnaseq v3.9 pipeline using STAR as aligner and Salmon for quantification with default parameters. Gene counts for each sample were log1p transformed, mean value among the two replicates was taken to compute Pearson correlation among gene expression pre- and post-time-lapse acquisition.

### Image Acquisition

Images were acquired with a Leica Thunder Imager (Leica Microsystems, Wetzlar, Germany), equipped with a Lumencor Spectra X Light Engine (Lumencor, Beaverton, USA) for fluorescence excitation, a motorized stage and a Leica DFC9000 GTC camera. For non-adherent cells, images were acquired with LAS X software (Leica Microsystems, Wetzlar, Germany, version 3.7.5.24914) using a 20X/0.75NA air objective and a binning 2×2 was applied to increase the SNR. The mCherry and mVenus signals were detected respectively with 540-580 nm and 460-500 nm excitation filters, 585 and 505 nm dichroic mirrors and 592-668 nm and 512-542 nm emission filters. The brightfield channel was also acquired for representation purposes. We imaged 20 to 25 fields of views per well and focal points were manually set in each position before starting the acquisition and kept constant during the whole time-lapse thanks to the Adaptive Focus Control (AFC, Leica Microsystems). The total duration of the time-lapse on non-adherent cells was 72 hours, and the time interval was set to 1 hour to prevent cell phototoxicity.

Regarding the experiment on adherent cells, a 10X/0.32 NA PH1 dry objective was used and the time-lapse duration was set to 120 hours, with 30 min as timestep. In this case 10 fields of view per well were acquired.

### Image Analysis

The image pre-processing step was performed using a custom-made Fiji macro. Briefly, the pipeline executed a flat-field correction on the fluorescence channels with the plugin Basic (Peng et al., 2017), then applied a Gaussian blur (sigma=1 px), a Top Hat filter (radius=20 px) and a background subtraction to enhance the cells’ signal; duplicated images of Red and Green channels were merged in a multichannel stack and brightness and contrast values were adjusted, according to each experiment, within the range of 0 to the maximum gray value of the stack’s histogram; the multichannel images were then converted into RGBs for each frame, and finally into a HSB stack from which the Brightness channel and the Hue channel were kept respectively for tracking and cell-phase profiling purposes; the final processed image was saved for further analysis and was composed of 4 channels: the Red and Green channels, the Brightness channel and the Hue channel.

The tracking analysis was realized through TrackMate (version 7.10.2). We automatized the execution of TrackMate over all the fields of view acquired in the experiment by adapting a Jython script, freely available on the website (https://imagej.net/plugins/trackmate/scripting/scripting). As the shape of our cells was approximately round, the Laplacian of Gaussian (LoG) detector was selected as the cell identifier on the Brightness channel, and the Linear Assignment Problem (LAP) Tracker algorithm was utilized for linking phase, adjusting the Max Linking distance parameter for each experiment. Gap closing was enabled for up to 3 frames, considering the significant decrease in fluorescence intensity during mitosis and the transition between G1 and S phase, that can eventually affect the detection of a cell, due to the rapid decrease of the Red signal and the relatively slow increase of the Green one (Sakaue-Sawano et al., 2017). Identified tracks were filtered for total duration, track displacement, and starting frame of the track. Specifically, the duration filter maximized the chances to follow at least one cell cycle (Track Duration > 25 hours), while the cutoff on total displacement assures the deletion of dead cells that are usually motionless (Track displacement > 10 px). Tracks that start after the first 10 hours were discarded to avoid tracker errors that may give rise to unreliable cell cycle quantification as the cells tend to form clusters above 24-30 hours of experiment (movie S1). The specific values of parameters of the LoG cells’ detector (i.e., spot radius, quality threshold) were preliminarily checked on images via the TrackMate GUI, as well as the linking and gap closing max distances and the final filters on tracks’ duration, displacement and starting frame. The parameters selected manually were then inserted in the script for the batch execution. As an output, we automatically saved a table of spots for each field of view, corresponding to cells in identified tracks, with the selected TrackMate features (See Suppl. Table I).

### Model Creation

To assess the traceability of each track in a fast, efficient and scalable manner, we employed a machine-learning approach. A dataset consisting of 2319 manually annotated tracks, with 1,939 tracks classified as untraceable and 380 tracks marked as traceable (~1/5) was used. The dataset was divided into training and test sets, with 1855 tracks (~ 80%) allocated for training and 464 tracks (~20%) reserved for testing while also stratifying for the outcome variable. To address the problem of the unbalanced outcome a downsampling procedure is employed in the training set.

Time-series-like features associated with each track are extracted (Wang, Smith and Hyndman, 2006; Hyndman et al., 2023), as illustrated in Fig. S3A, displaying the distribution of the 70 features utilized for distinguishing between traceable and untraceable cells.

To optimize model performance and address potential biases of the random forest model employed, we applied stratified 10-fold cross-validation on the training set, creating resampling folds. Leveraging these folds, we conducted hyperparameter tuning. Specifically, we explored different values for Randomly Selected Predictors (mtry), number of trees (trees), and Minimal Node Size (min_n) to identify the combination that maximized the Area under the receiver operator curve (AUC) (Sacks et al., 1989). Subsequently, an additional round of training was conducted using the selected hyperparameters, this time employing the entire training set. The trained model was then evaluated on an independent test set to assess its performance. The results, as depicted in Fig. S3B, revealed an AUC value of 0.971 as well as a sensitivity of 0.897, a specificity of 0.974 and an accuracy of 0.905 confirming the validity of the training procedure. These values were comparable to those obtained from the resampling folds, indicating the absence of overfitting.

Given the relatively small size of the manually annotated training set, we performed an additional and final round of training using the pseudo-labelling framework (Lee, 2013). The original model was used to predict unlabeled data, any predicted probability exceeding 0.5 was considered indicative of confidence in the prediction. A new semi-supervised random forest model was trained on this augmented dataset, employing the same hyperparameters as determined previously. This final model achieved an improved AUC of 0.975 on the test set, indicating enhanced track trackability assessment. The schematics of the pipeline, implemented in R (v. 4.3.0) (R Core Team, 2021) using Tidymodels (v. 1.1.0) (Kuhn and Wickham, 2020), and Tidyverse (v.2.0.0) (Wickham et al., 2019) can be seen in Fig. 3 (panels A).

### Time series analysis

To generate time-series-like information, we utilized the above-mentioned Fiji pipeline, which provided a table containing intensity values for the red, green, and hue channels. To address missing frames resulting from tracking gaps, an Exponential Weighted Moving Average (EWMA) was fitted to impute these frames.

To further refine the track curves, we applied a two-step smoothing process. Initially, a Simple Moving Average (SMA) was applied, followed by fitting a fixed lambda Smoothing Spline with λ= 0.0001. Finally, min-max scaling is used to normalize the tracks within the range of [0, 1] to make the red and green intensity comparable.

To tackle tracking errors introduced by the Fiji pipeline, we applied the aforementioned random forest model to assess the traceability of each cell (Fig. 2B).

On the tracks classified as trackable, a manual threshold on the hue intensity is employed to determine the cell cycle phases (G1, S, and G2/M). Furthermore, to quantify the cell cycle at the single-cell level, track splitting to isolate individual cell components and perform phase assignment into G1, G1/S, S, and G2/M is performed using a custom R function.

The resulting single-cell tracks enabled the quantification of individual phase durations for each cell. As an example, the cell phase quantification for five different conditions can be observed in Fig. 3.

### Model Deployment

The resulting model was saved as a .rds file to facilitate practical implementation. Moreover, we encapsulated the model as an API in a Docker container using Vetiver (v. 0.2.1) (Silge, 2023), enabling easy deployment and usage.

### EdU incorporation and assessment with flow cytometry and imaging

A two-hour EdU pulse was performed by replacing half of the total media volume of the cells with 2X concentrated EdU in the corresponding growth medium, followed by subsequent fixation. Click-iT™ EdU Alexa Fluor™ 647 Flow Cytometry Assay Kit (CN: C10419, Thermo-Fisher Scientific, Waltham, MA, USA) and Click-iT™ EdU Cell Proliferation Kit for Imaging, Alexa Fluor™ 647 dye (CN: C10340, Thermo-Fisher Scientific, Waltham, MA, USA) were used for flow cytometry and imaging, respectively. The experiments were performed according to the manufacturer’s protocols for the mentioned kits.

DNA staining with DAPI using 500 μL of 5 μg/mL DAPI in PBS for 10^6^ cells, followed by overnight incubation at 4°C, was additionally performed for cell cycle profiling by flow cytometry. Alternatively, 5 μg/mL Hoechst® 33342 (Thermo-Fisher Scientific, Waltham, MA, USA) was used to stain DNA for imaging purposes.

### Confocal imaging and image analysis

The confocal images were captured using the Eclipse Ti2 microscope (Nikon Europe B.V.) combined with the X-Light V3 spinning disk (CrestOptics S.p.A.), solid-state lasers from the Lumencor Celesta light engine, a multiband dichroic mirror, single-band emission filters, and an sCMOS camera (Kinetix, Teledyne Photometrics). A total of 144 and 464 fields of view (FOV) were acquired using a PLAN APO λD 60x 1.42/NA oil immersion objective lens (pixel size of 116×116 nm) for the Alexa647 (labeling EdU), mCherry, mVenus, and DAPI signals in Kasumi-1 and MDA-MB-231 cells, respectively.

To assess the mean intensity of each signal within the nuclei, a custom Python-based image segmentation pipeline utilizing the Stardist deep learning algorithm (Schmidt et al., 2018) was employed. The Hoechst® 33342 intensity corresponding to the 2N DNA amount was calculated, and cells in the initial S phase were selected based on being EdU positive, having mVenus and mCherry intensities above background, and a DNA content smaller than 1.15 * 2N. All data analysis was conducted using RStudio software.

## Code availability

The source code and user manual for the Fiji pipeline is available at https://github.com/ieoresearch/cellcycle-image-analysis.

The source code and user manual for the R pipeline is available at GitHub repository.

## Funding and Acknowledgements

This work was partially supported by the Italian Ministry of Health with Ricerca Corrente and 5×1000 funds and by the Grant AIRC IG20 to SM. KH was supported by Marie Skłodowska-Curie Innovative Training Network (grant no. 813327 ‘ChromDesign’) and AIRC fellowships.

The authors are thankful to Dr. Marina Mapelli and Michela Bruzzi for their kind help in performing EdU experiments; and to Dr. Mattia Marenda and Giulia Tini and for their kind assistance in image and data analysis.

## Supplementary Figure legends

**Figure S1:**
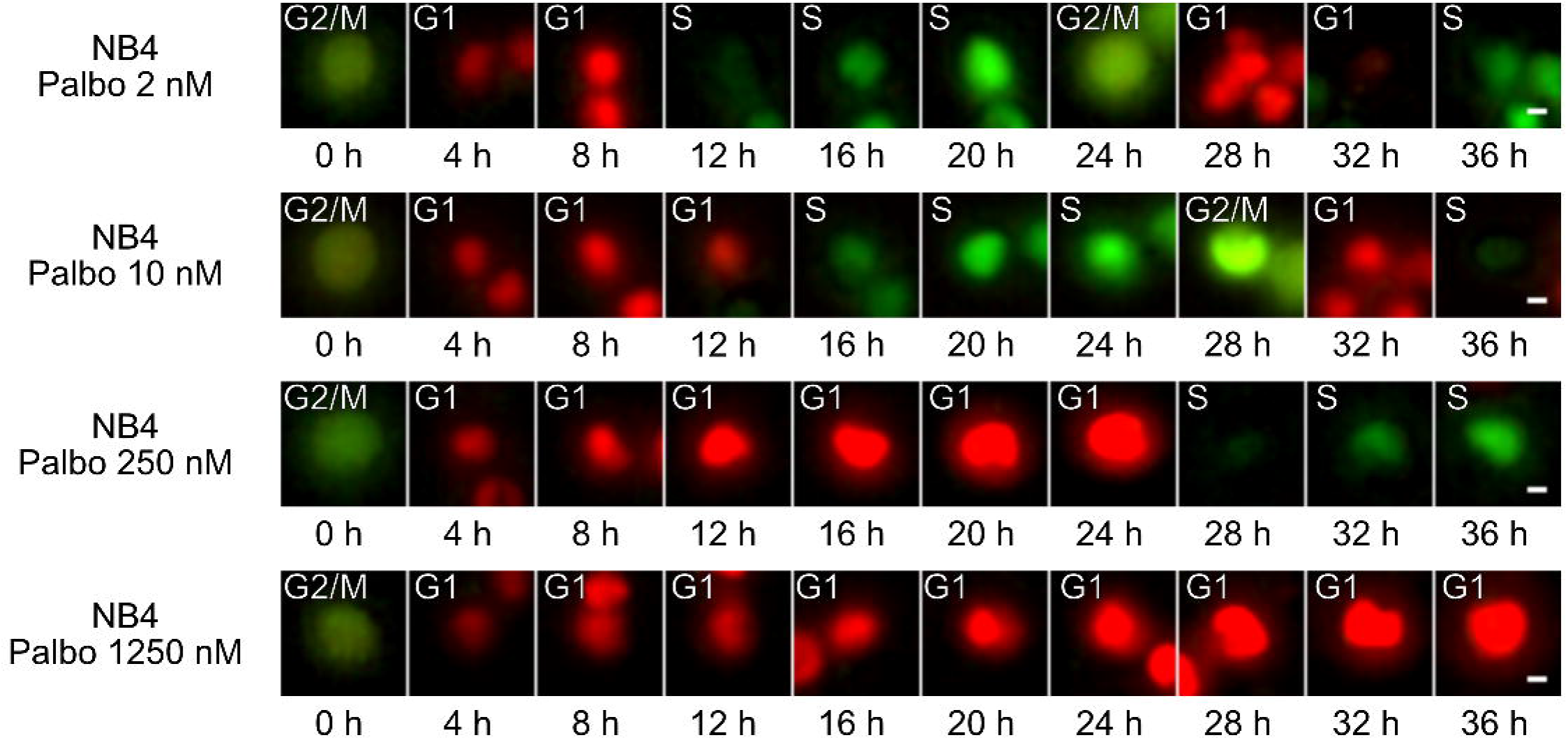
Cell Cycle Phases in Treated NB4 cells. A) Examples from NB4 cell line upon different Palbociclib treatment concentrations showing a cell that explores the cell cycle phases (Scale bar is 5 µm). For Palbociclib 250 nM and Palbociclib 1250 nM conditions, the cell was manually tracked for presentation purposes.

**Figure S2:**
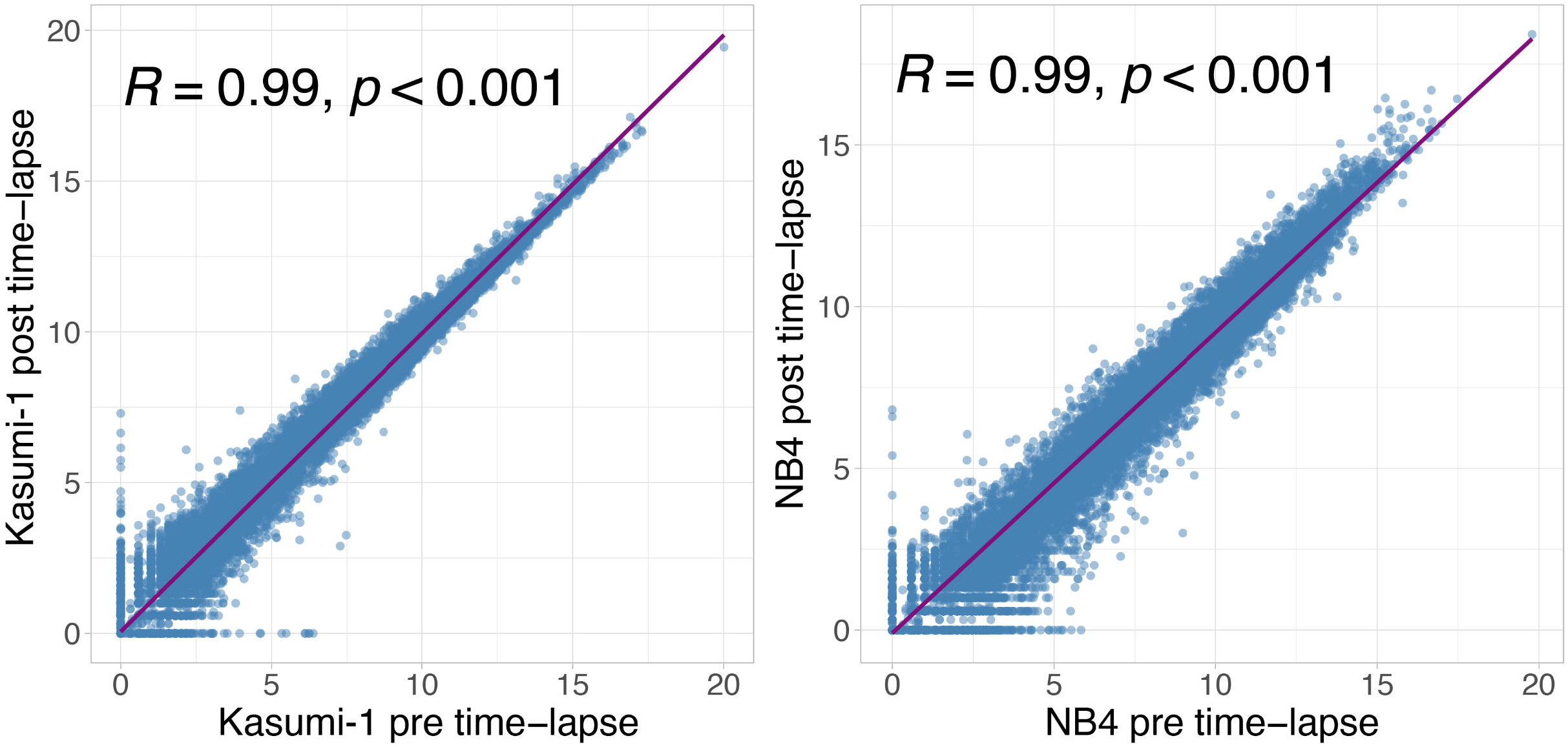
Transcriptome correlation analysis of AML cells seeded on SBS. A) Pearson correlation among pre and post time-lapse acquisition of Kasumi-1 (left) and NB4 (right) cell lines. In the x axis is reported the log1p read counts mean between the two replicates before the acquisition, while on the y axis the log1p read counts mean between the two replicates after the acquisition.

**Figure S3:**
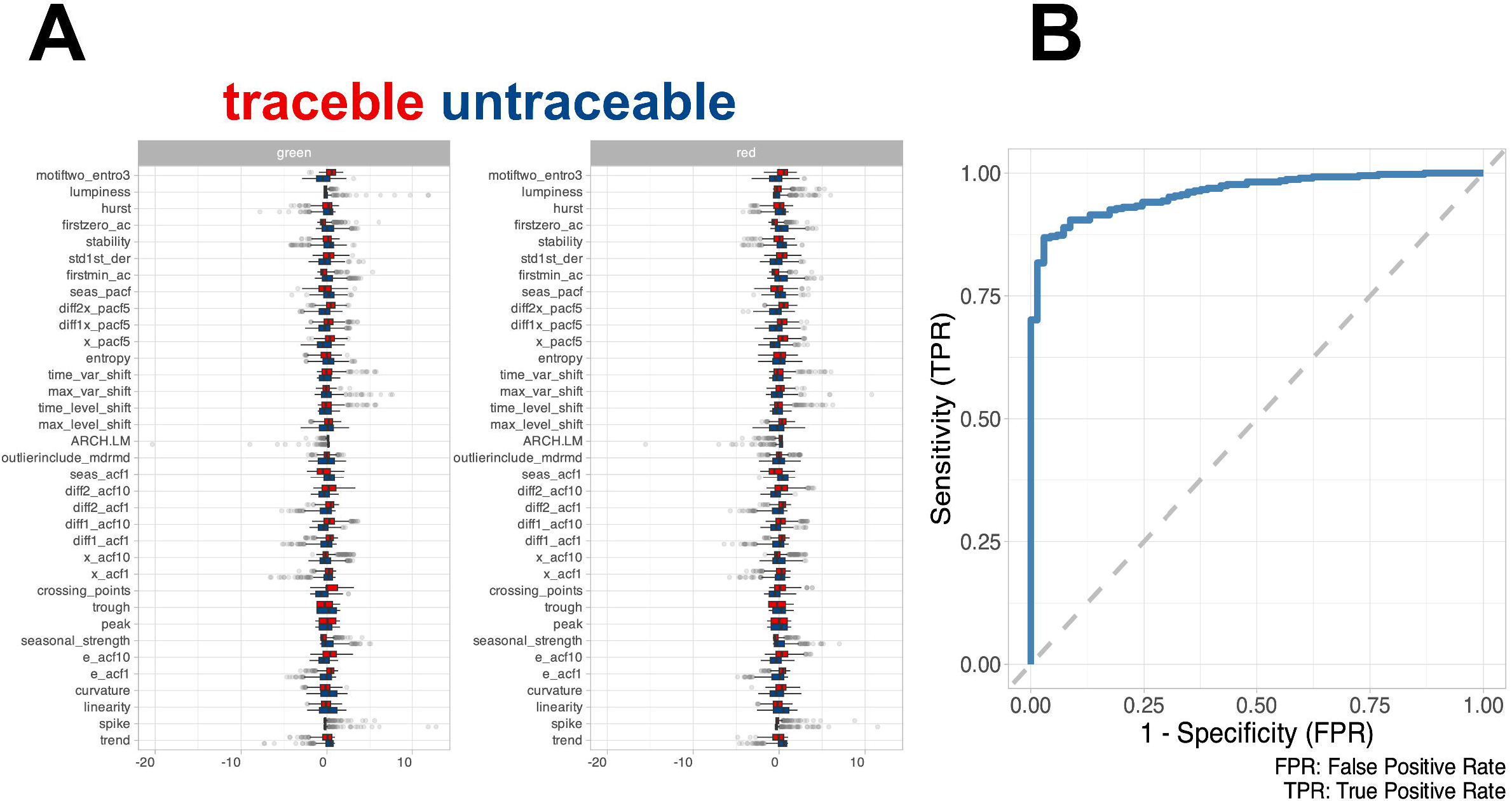
Feature Distribution and Model Performance. A) Boxplot showing the distribution of the feature extracted from the training set using timetk. B) ROC curve of the random forest model computed in the test set.

**Figure S4:**
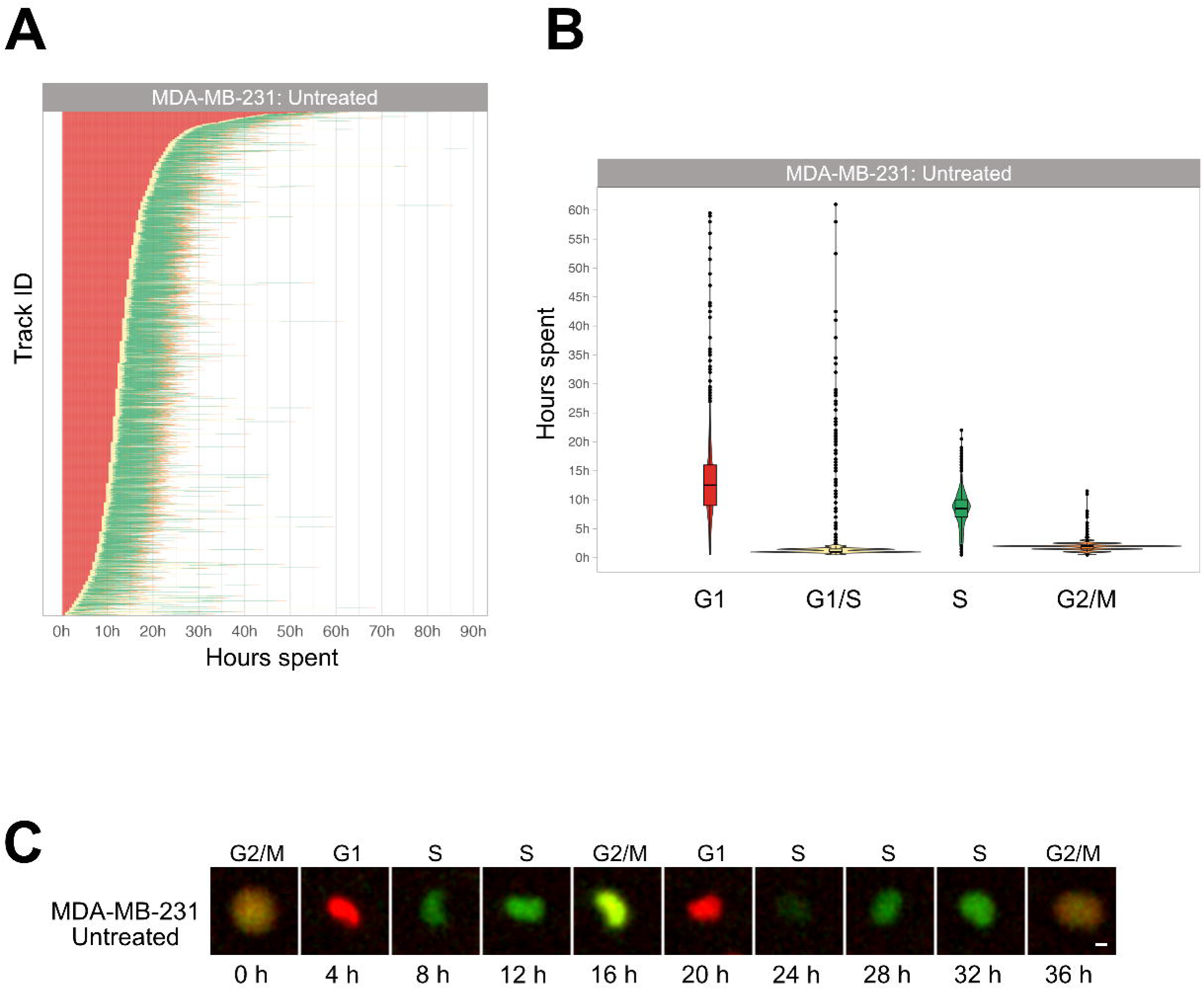
Cell Cycle Phase Assignment in MDA-MB-231 cells. A) Waterfall plot of untreated MDA-MB-231 cells. Each row corresponds to a single cell. B) Boxplot of the cell phase duration of the first cell cycle, as obtained by the pipeline, without excluding of the outliers. A total of 1116 cells were analyzed, obtaining a mean (± SD) cell cycle duration of 24.5 ± 8.5 h. C) Example of an adherent cell in control condition tracked with the described pipeline (Scale bar is 5 µm).

**Figure S5:**
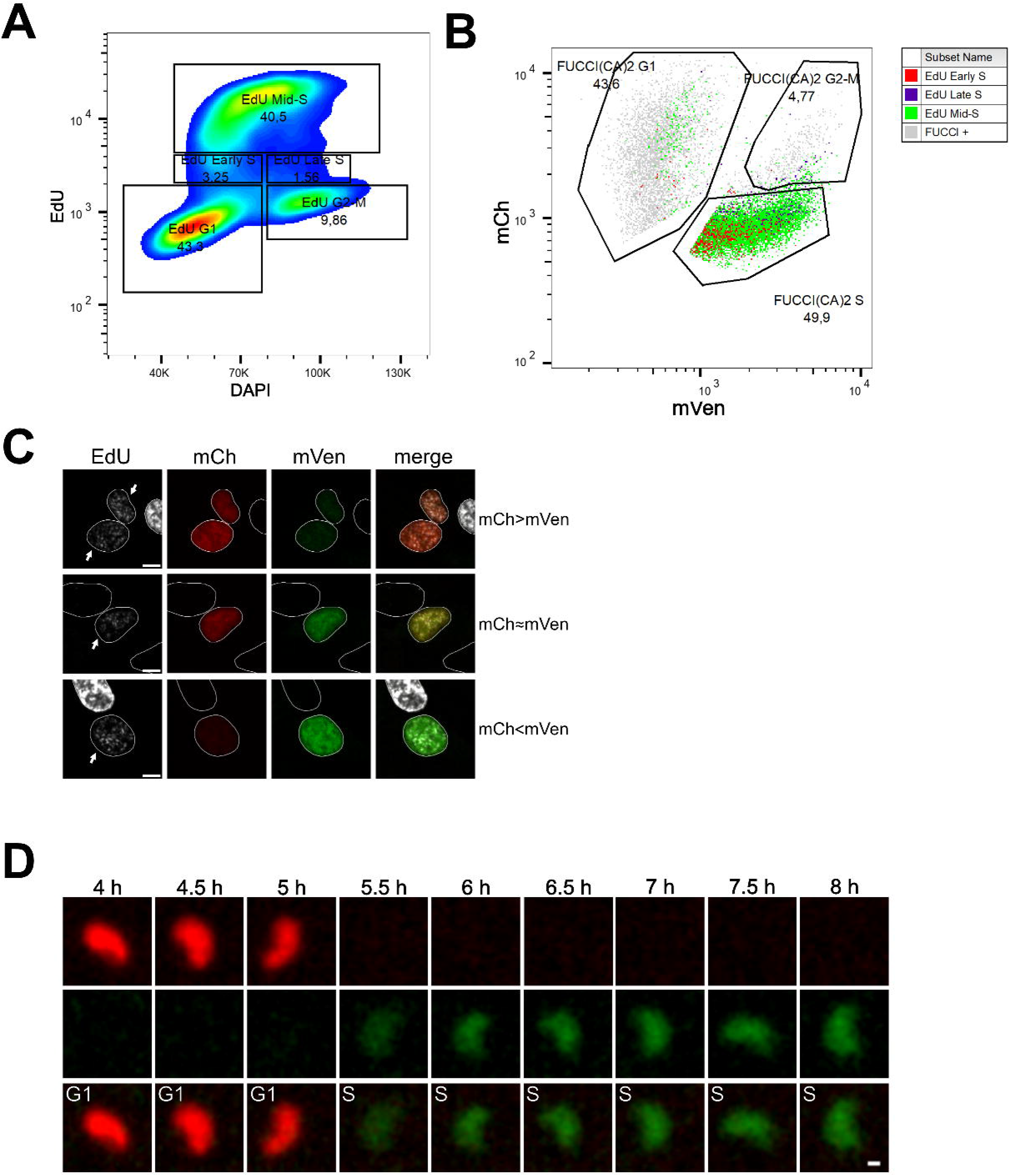
Cell Cycle Profiling and G1 to S Transition Analysis in MDA-MB-231 cells. A) Density plot showing the cell cycle profiling of FUCCI(CA)2-MDA-MB-231 cells based on EdU vs DAPI. B) Scatter plot of the same cells in “A” showing the cell cycle profiling based on FUCCI(CA)2. Cells identified to be in early, mid, and late S phase by EdU are highlighted in red, green, and purple, respectively. Due to the chosen gates and the 2-hour EdU pulse, a small percentage of EdU-positive cells leaked into the G1 and G2-M FUCCI(CA)2-defined groups. C) Representative images of nuclei in the G1 to S transition are indicated by arrows. Depending on the exact time after S phase initiation, the overlay of the mCh and mVen channels can show a red prevalence (first row), a yellow color (middle row), or a green prevalence (third row). Scale bar is 5 µm. D) The frames of the untreated MDA-MB-231 cell shown in Fig. S4C, are presented from the 4h to the 8h frame. The scale bar represents 2 µm.

**Movie S1:** Representative movie of NB4 cells seeded on SBS + MC.

**Movies S2-S4:** Representative movies of 3 cells tracked with TrackMate expressing the FUCCI(CA)2 probe, showing the sequence of color changes (1 frame/hour).

**Movie S5:** Representative movie of NB4 cells seeded on SBS without MC.

